# The HSV-1 immediate early protein ICP22 interacts with the human antisense function 1 protein to promote viral replication

**DOI:** 10.64898/2026.06.24.734377

**Authors:** Yu Ye, Zhijun Yang, Mengzhou Xue, Chunfu Zheng

**Affiliations:** College of Animal Science and Technology, Jiangxi Engineering Research Center for Animal Health Products, Jiangxi Agricultural University, Nanchang, 330045, Jiangxi, China; Shanghai Omics Biotechnology Co., Ltd.; Department of Cerebrovascular Diseases, The Second Affiliated Hospital of Zhengzhou University, Zhengzhou, China, 450001; Department of Microbiology, Immunology and Infectious Diseases, University of Calgary, Calgary, Alberta, Canada

**Keywords:** HSV-1, ICP22, ASF1, histone H3

## Abstract

Herpes simplex virus type 1 (HSV-1) is a common human pathogen that undergoes lytic replication in epithelial and other permissive cell types and can establish latency in peripheral neurons. ICP22 is a multifunctional HSV-1 immediate-early protein that localizes to the nucleus of infected cells; however, its interactions with host cellular factors remain incompletely understood. Here, ICP22 was demonstrated to interact with the human antisense function 1 protein (ASF1), including both ASF1a and ASF1b, in transfected cells and HSV-1-infected cells, respectively. ICP22 also colocalized with ASF1 in the nucleus. ICP22 amino acids 213 to 340 are important for the interaction of ICP22 with ASF1, whereas amino acids 37 to 153 of ASF1a and ASF1b are critical for their interactions with ICP22. Furthermore, ICP22 expression was associated with reduced ASF1-H3.1 co-immunoprecipitation under the tested conditions. ASF1 knockdown also reduced HSV-1-BAC-Luc luciferase output, indicating that ASF1 contributes to efficient infection-associated reporter activity in this study. Collectively, these results indicate that the interaction of HSV-1 ICP22 with ASF1 might help regulate the transcription of viral or cellular genes during HSV-1 infection.

## Introduction

Herpes simplex virus 1 (HSV-1) is a common and widely studied human pathogen that can replicate in epithelial cells and other cells of the host or can remain latent in peripheral neurons. During productive infection, the 152-kb double-stranded HSV-1 genome is rapidly translocated to the nucleus, where the viral genes are transcribed by the host cell RNA polymerase II (Pol II) in a temporally orchestrated program [1–6]. The expression of the ∼ 80 viral genes occurs in a coordinately activated cascade that consists of the sequential expression of immediate-early (IE), early (E), and late (L) genes.

The transcription of IE genes by the host cell RNA Pol II is stimulated by the virion tegument protein VP16 and does not require prior viral protein synthesis. This results in the synthesis of five infected cell proteins (ICPs) [7]. Two of these genes, ICP4 and ICP27, are essential for HSV-1 replication [8]. The interaction between ICP22 and facilitates chromatin transcription (FACT) is important for HSV-1 gene expression and pathogenicity [7]. In contrast, the ICP0, ICP22, and ICP47 proteins appear to be dispensable in continuously growing cell cultures and are therefore considered to be nonessential [9]. However, it has also been reported that ICP22 is required for optimal virus replication in primary human cells and in cell lines of rodent origin, indicating that the regulatory protein ICP22 has some cell type-dependent activity [10].

ICP22 consists of 420 residues and is encoded by a spliced mRNA transcribed from the US1 gene. It is necessary for efficient HSV-1 growth in animal models of infection [11] as well as for efficient in vitro growth in some, but not all, cultured cells [12]. For instance, human embryonic lung (HEL) cells do not support the growth of ICP22 mutants, while African green monkey kidney (Vero) cells do. During infection, UL13 and another viral protein kinase, US3, are the main phosphorylators of ICP22 [13]. In addition to inducing the modification of the host cell RNA Pol II, several other functions have been attributed to ICP22 [14, 15], including the induction of certain viral L genes, the alteration of cell cycle-related proteins [16], and the determination of virion composition [11]. It is clear that ICP22 is a multifunctional protein localized to the nucleus of infected cells [17, 18]; however, the host cellular factors of ICP22, as well as the biological functions of their interactions, are still poorly understood.

Virus-host protein interactions are critical for successful herpesvirus infection. Previous studies have shown that ASF1 contributes to herpesvirus gene expression and replication[19] and that the varicella-zoster virus IE63 protein, a homolog of HSV-1 ICP22, interacts with ASF1 [20]. However, whether HSV-1 ICP22 physically associates with ASF1 and how this interaction may influence ASF1-related chromatin functions remains unclear. In this study, we examined the interaction between HSV-1 ICP22 and human ASF1, mapped the regions involved, and evaluated its potential functional relevance in cultured-cell infection models.

## MATERIALS AND METHODS

### Cells and viruses

HEK293T, HeLa, and Vero cells were maintained in Dulbecco’s modified Eagle’s medium (Invitrogen) supplemented with 10% FBS, penicillin, streptomycin, and glutamine. Wild-type HSV-1 (F strain) and recombinant HSV-1-BAC-Luc have been reported previously [21]. All viruses were propagated and titrated in Vero cells.

### Plasmids and Antibodies

Plasmids expressing myc-tagged human ASF1a (p408) and ASF1b (p542) under the simian virus 40 and T7 promoters were kind gifts from Dr. P. D. Adams as described previously [22]. The PCR products corresponding to amino acids (aa) 1-153, 37-204 of ASF1a and 1-153, 37-202 of ASF1b were amplified from the Myc-ASF1a and Myc-ASF1b plasmids via primers containing *Eco*RI and *Bam*HI restriction enzyme sites and subsequently cloned and inserted into the pCMV-Myc vector to generate pMyc-ASF1a-153, pMyc-ASF1a-37, pMyc-ASF1b-153 and pMyc-ASF1b-37, respectively. The plasmids pcDNA22 and pcDNAUS1.5 encode N-terminally Flag-tagged ICP22 and US1.5 (assuming that US1.5 corresponds to residues 147 to 420 of ICP22), respectively, and pcDNA22BA and pcDNA22PS were kindly provided by Dr. Stephen A. Rice as described previously [23]. The pICP4-Luc and pTK-Luc reporters were used as previously described [24]. The anti-Flag and anti-Myc monoclonal antibodies (mAbs) (Santa Cruz) and rabbit anti-ICP22 polyclonal antibody (pAb) were kind gifts from Dr. Bernard Roizman, and the anti-ASF1 pAb was purchased from Santa Cruz.

### Co-immunoprecipitation and Western blot analysis

Co-immunoprecipitation (Co-IP) and Western blotting were performed as described in our previous studies [24, 25]. The cell lysates and immunoprecipitated proteins were separated on denaturing 12% polyacrylamide gels and transferred to nitrocellulose membranes. The transferred proteins were probed with anti-Myc and anti-Flag mAbs at a 1:1000 dilution in PBS.

### Transfection and fluorescence microscopy

Transfection and fluorescence microscopy experiments were performed as previously described [25, 26]. All photomicrographs were taken at a magnification of 400× unless otherwise specified. Each photomicrograph represents a vast majority of the cells with similar subcellular localizations.

### Reporter gene assays

Luciferase assays were performed to measure viral promoter activity. HEK293T cells were cotransfected with 200 ng of pICP4-Luc or pTK-Luc and 0.5 µg of the plasmid ICP22-Flag or the plasmid Myc-ASF1a. Twenty-four hours after transfection, luciferase expression was detected via a luciferase assay kit (Promega) according to the manufacturer’s manual.**siRNA-mediated knockdown and HSV-1-BAC-Luc infection**

HEK293T cells were transfected with siRNA oligos against ASF1a or ASF1b via the Lipofectamine 2000 reagent as suggested by the manufacturer (Invitrogen). The 19-nucleotide siRNAs against human ASF1a (siRNA,

5’-AAGUGAAGAAUACGAUCAAGU-3’) and ASF1b (siRNA, 5’-AACAACGAGUACCUCAACCCU-3’) were obtained from RiboBio. At 24 h after siRNA transfection, the cells were infected with HSV-1-BAC-Luc virus at an MOI of 5, and the luciferase activities were determined to measure the viral replication levels at 16 h post infection.

## RESULTS

### HSV-1 ICP22 associates with human ASF1 in transfected and HSV-1-infected cells

A previous study reported that the varicella-zoster virus (VZV) immediate-early 63 protein (IE63) could interact with the human ASF1 protein [20]. To investigate whether HSV-1 ICP22 interacts with ASF1, 293T cells were cotransfected with pcDNA-ICP22 and pMyc-ASF1a or pMyc-ASF1b for co-IP analysis. As a result, ICP22-Flag was successfully immunoprecipitated by Myc-ASF1a or Myc-ASF1b using an anti-Myc mAb but not with nonspecific mouse IgG (Fig. 1A), suggesting that ICP22 interacted with ASF1a and ASF1b, respectively. To determine whether these interactions were present in HSV-1-infected cells, 293T cells were mock-infected or infected with HSV-1(F) virus at a multiplicity of infection (MOI) of 5 PFU per cell. Twenty hours after infection, cell lysates were prepared and subjected to co-IP analysis with an anti-ASF1 pAb followed by immunoblotting to detect the ICP22 and ASF1 proteins. As expected, ICP22 coimmunoprecipitated with both the ASF1a and ASF1b proteins in HSV-1-infected cells but not in mock-infected cells (Fig. 1B). Overall, ICP22 was shown to interact with both ASF1a and ASF1b in both overexpressing and HSV-1-infected cells.

**Fig. 1.**
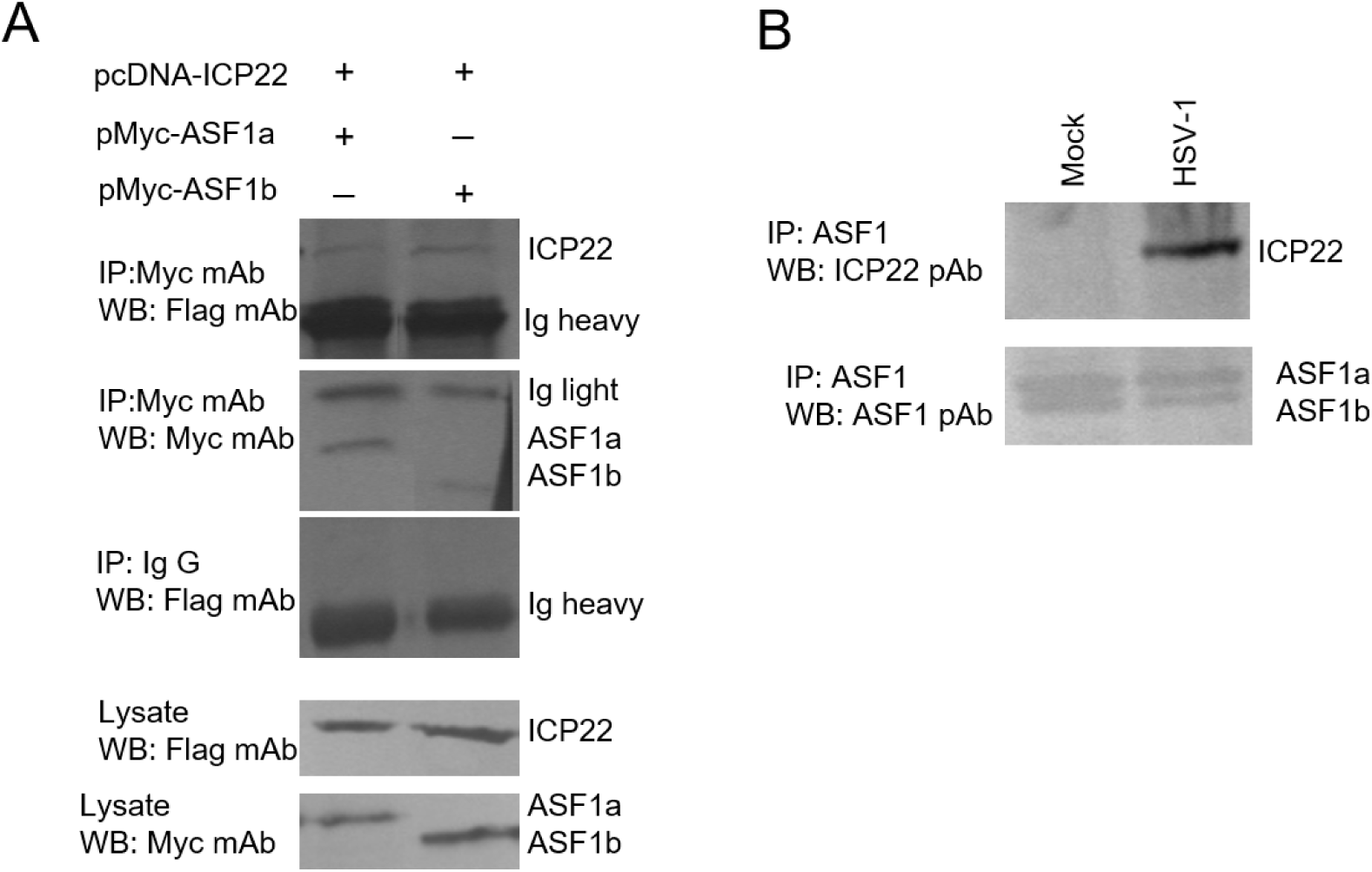
Interaction of HSV-1 ICP22 with human antisense function 1 protein (ASF1) in transfected and HSV-1-infected cells. (A) HEK293T cells were cotransfected with pcDNA-ICP22 and pMyc-ASF1a or pMyc-ASF1b. At 36 h after transfection, cells were lysed and subjected to immunoprecipitation with anti-Myc antibody or matched IgG control, followed by immunoblotting with anti-Myc and anti-Flag antibodies. (B) HEK293T cells were mock-infected or infected with wild-type (WT) HSV-1 at an MOI of 5 for 16 h. Cell lysates were immunoprecipitated with anti-ASF1 antibody and analyzed by immunoblotting for ASF1 and ICP22.

### The amino acids 213 to 340 of HSV-1 ICP22 interact with the highly conserved domain of both ASF1a and ASF1b

HSV-1 ICP22, a highly conserved protein in alphaherpesvirus, is a multifunctional regulatory protein localized to the nucleus of infected cells [27]. To determine which domain of ICP22 affects its interaction with ASF1, a series of ICP22 deletion mutants, including pcDNAUS1.5, pcDNA22-BA and pcDNA22-PS, were employed to investigate their interactions with ASF1 via co-IP analysis (Fig. 2A). As a result, US1.5-Flag and ICP22-BA were immunoprecipitated with ASF1a and ASF1b, respectively, via an anti-Myc mAb (Fig. 2A) but not with nonspecific mouse IgG (data not shown). However, neither ASF1a nor ASF1b was immunoprecipitated with ICP22-PS via an anti-Myc mAb (Fig. 2A). Therefore, amino acids (aa) 213 to 340 of ICP22 are important for its interaction with both ASF1a and ASF1b.

**Fig. 2.**
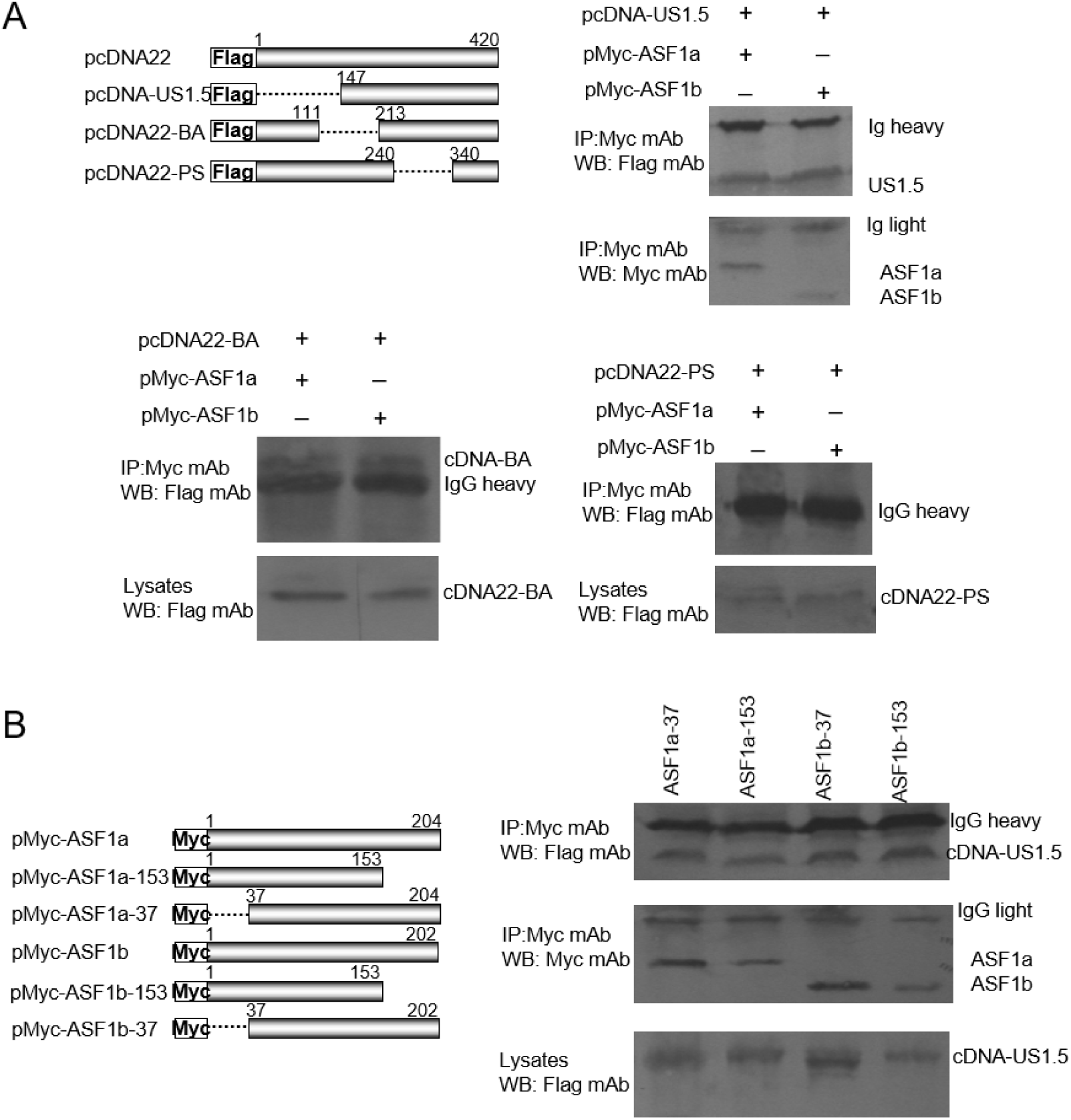
A central region of ICP22 and the conserved core of ASF1 contribute to the ICP22–ASF1 interaction. (A) HEK293T cells were cotransfected with pMyc-ASF1a or pMyc-ASF1b and the indicated ICP22 constructs. At 36 h after transfection, lysates were subjected to immunoprecipitation and immunoblotting as described in Fig. 1A. (B) HEK293T cells were cotransfected with pcDNA-US1.5 and pMyc-ASF1a-153, pMyc-ASF1a-37, pMyc-ASF1b-153, or pMyc-ASF1b-37 plasmids. At 36 h after transfection, lysates were analyzed by co-immunoprecipitation and immunoblotting.

Human ASF1a and ASF1b share 71% amino acid identity. The most conserved area of the two proteins, between amino acids 37 and 155, shares 90% identity, whereas amino acids 1 to 30 share 70% identity. Next, we tested which domains of ASF1a and ASF1b are important for their interactions with HSV-1 ICP22. Both Myc-ASF1a-153 and Myc-ASF1a-37 were immunoprecipitated with US1.5-Flag via an anti-Flag mAb (Fig. 2B), suggesting that the highly conserved aa 37 to 153 of ASF1a are very important for its interaction with ICP22. Similarly, both Myc-ASF1a-153 and Myc-ASF1a-37 coimmunoprecipitated with US1.5-Flag (Fig. 2B). Collectively, the conserved domains (aa 37 to 153) of both ASF1a and ASF1b were shown to be important for their interactions with ICP22.

### ICP22 colocalizes with ASF1 in HSV-1-infected cells

ICP22 is located in the nucleus during lytic replication of HSV-1[25]. ASF1 localizes predominantly to the nuclei of cells [28]. To test whether ICP22 and ASF1 colocalize in the nucleus during HSV-1 lytic infection, HeLa cells were cotransfected with the plasmids ICP22-Flag and Myc-ASF1a or Myc-ASF1b, respectively. As expected, ICP22 colocalized with both ASF1a and ASF1b in the nucleus (Fig. 3). Thus, the colocalization of ASF1a and ASF1b with ICP22 in the nucleus of infected cells further supports their interaction.

**Fig. 3.**
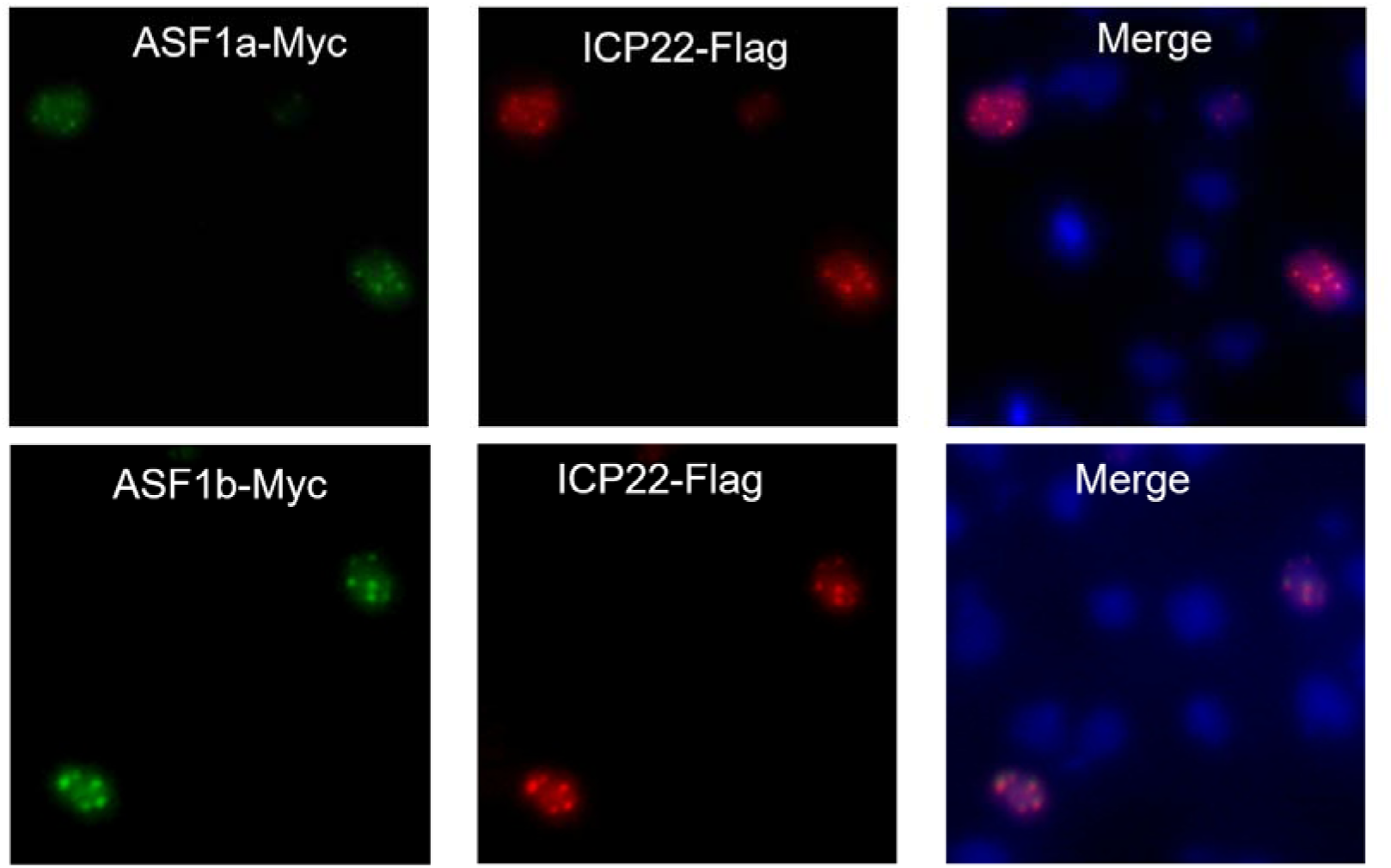
Subcellular colocalization of HSV-1 ICP22 with both ASF1a and ASF1b. HeLa cells were cotransfected with the pcDNA-ICP22 and pMyc-ASF1a or pMyc-ASF1b plasmids. At 24 h posttransfection, the cells were fixed, permeabilized, and stained with mouse anti-myc mAb and rabbit anti-Flag pAb, followed by incubation with FITC-conjugated goat anti-mouse (green) and TRITC-conjugated goat anti-rabbit (red) IgG antibodies. The cell nuclei (blue) were stained with Hoechst 33258. The images were obtained via fluorescence microscopy using a ×40 objective.

### ASF1 abrogates the repression of viral promoter activity by ICP22

Previous studies have suggested that ICP22 can repress transcription from a large number of promoters in transient reporter assays. To investigate whether the interaction between ASF1 and ICP22 affects the repression of viral promoter activity by ICP22, 293T cells were cotransfected with the pICP4-Luc or pTK-Luc plasmid and the plasmids ICP22-Flag and Myc-ASF1a. Twenty-four hours post-transfection, total cell lysates were prepared, and luciferase activities were measured. Transfection of the ICP4 or TK reporter plasmid alone resulted in high levels of firefly luciferase expression (Fig. 4), whereas cotransfection of ICP22 significantly repressed the luciferase promoter activity of ICP4-Luc or TK-Luc (Fig. 4). However, the coexpression of ASF1a abrogated the repression of ICP4 and TK promoter activity by ICP22 (Fig. 4). Overall, ASF1 was shown to abrogate the repression of viral promoter activities by ICP22.

**Fig. 4.**
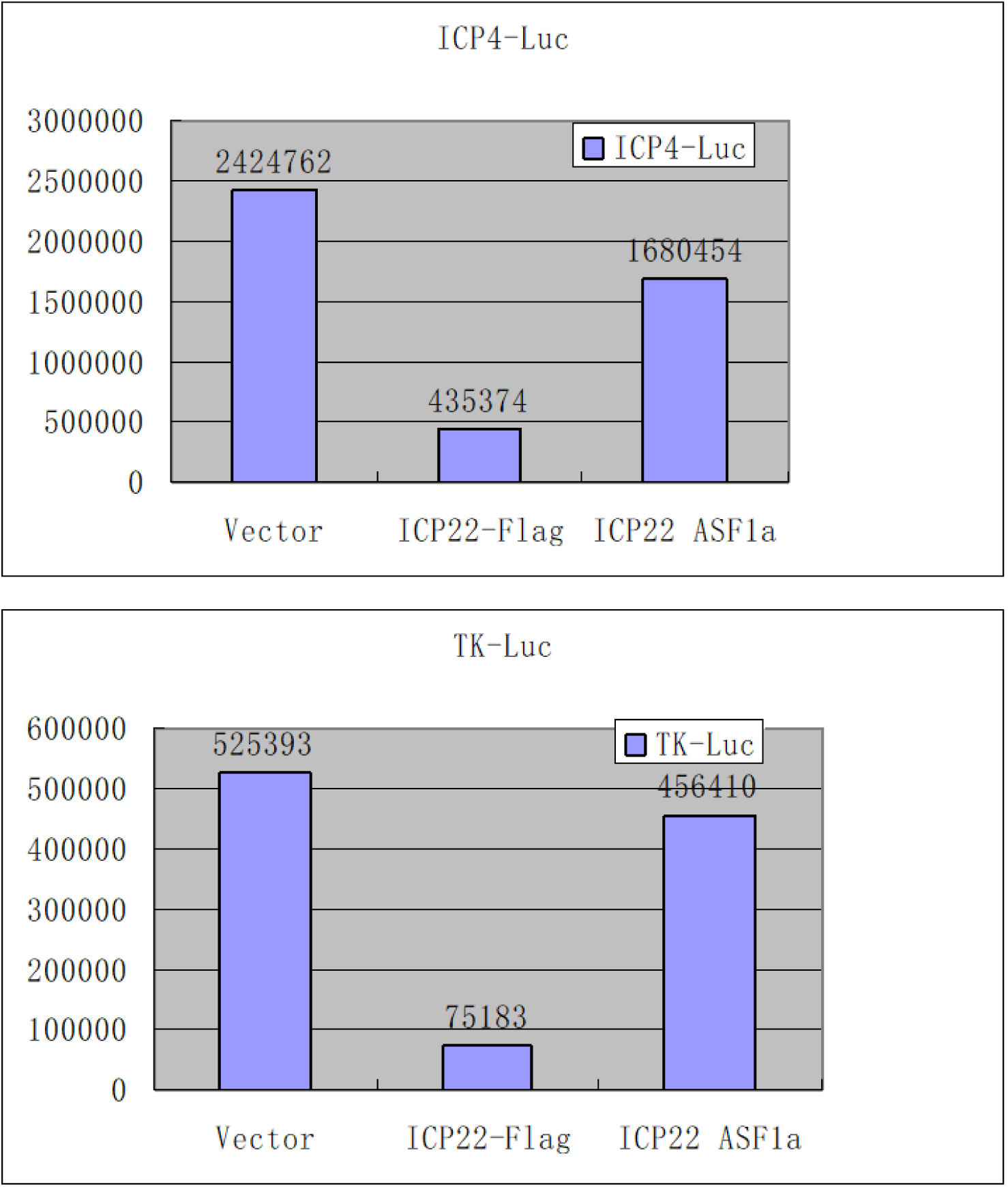
Ectopic expression of ASF1a attenuates the repression of viral promoter activity by ICP22. HEK293T cells were cotransfected with 200 ng of pICP4-Luc or pTK-Luc and 0.5 µg of the plasmid ICP22-Flag or the plasmid Myc-ASF1a. 24 h after transfection, luciferase expression was detected via a luciferase assay kit. The data presented are the averages of the results of three experiments.

### Both ASF1a and ASF1b contribute to HSV-1-BAC-Luc reporter output

To the contribution of ASF1 to HSV-1 infection-associated reporter output, 293T cells were transfected with both ASF1a- and ASF1b-specific siRNAs. The results revealed that the knockdown of either ASF1a or ASF1b reduced HSV-1-BAC-Luc replication (approximately 2-fold) (Fig. 5). However, combined knockdown produced a stronger reduction (Fig. 5). These findings indicate that both ASF1 isoforms contribute to efficient HSV-1-BAC-Luc reporter output under the conditions tested. Because this assay measures infection-associated luciferase signal rather than viral DNA replication directly, the data are interpreted accordingly.

**Fig. 5.**
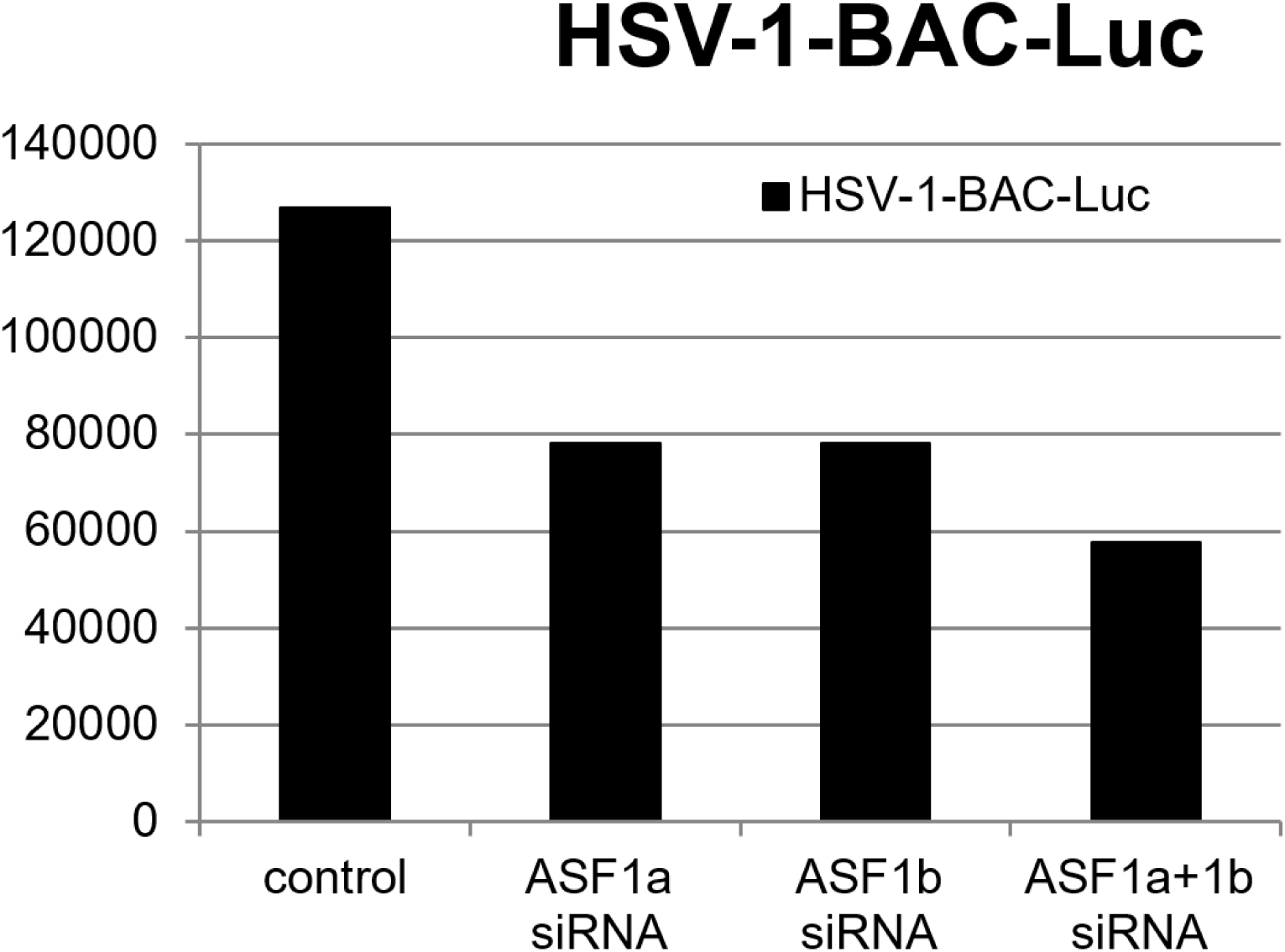
Both ASF1a and ASF1b contribute to HSV-1-BAC-Luc.infection-associated luciferase output. HEK293T cells were transfected with siRNA oligos against ASF1a or ASF1b via the Lipofectamine 2000 reagent. At 24 h after siRNA transfection, the cells were infected with HSV-1-BAC-Luc virus at an MOI of 5, and the luciferase activities were determined to measure the viral replication levels at 16 h post infection.

### Overexpression of ICP22 abrogates the interaction between H3.1 and ASF1a

ASF1 interacts with histones H3 and H4 and plays a major role in nucleosome assembly and disassembly during DNA replication, transcription, and DNA repair [29]. Human ASF1a and ASF1b interact with histone H3 isotypes H3.1 and H3.3 [30]. To determine whether the interaction between ICP22 and ASF1 interferes with the interaction between ASF1 and H3, 293T cells were transfected with the plasmids ASF1a-Myc and/or ICP22-Flag. The cell lysates were immunoprecipitated with anti-H3.1 pAb, followed by immunoblotting with anti-Flag and anti-Myc mAbs. As a result, in cells expressing ASF1a-Myc, ASF1a was found to be coimmunoprecipitated with H3.1 by the anti-H3.1 pAb (Fig. 6) but not with nonspecific rabbit IgG (data not shown). However, ICP22 expression abrogated the interaction between ASF1a and H3.1 via the anti-H3.1 pAb (Fig. 6). Overall, the overexpression of ICP22 abrogated the interaction between ASF1 and H3.1, suggesting that the interaction between ASF1 and H3.1 may play an important role in the regulatory functions of ICP22 in HSV-1 infection.

**Fig. 6.**
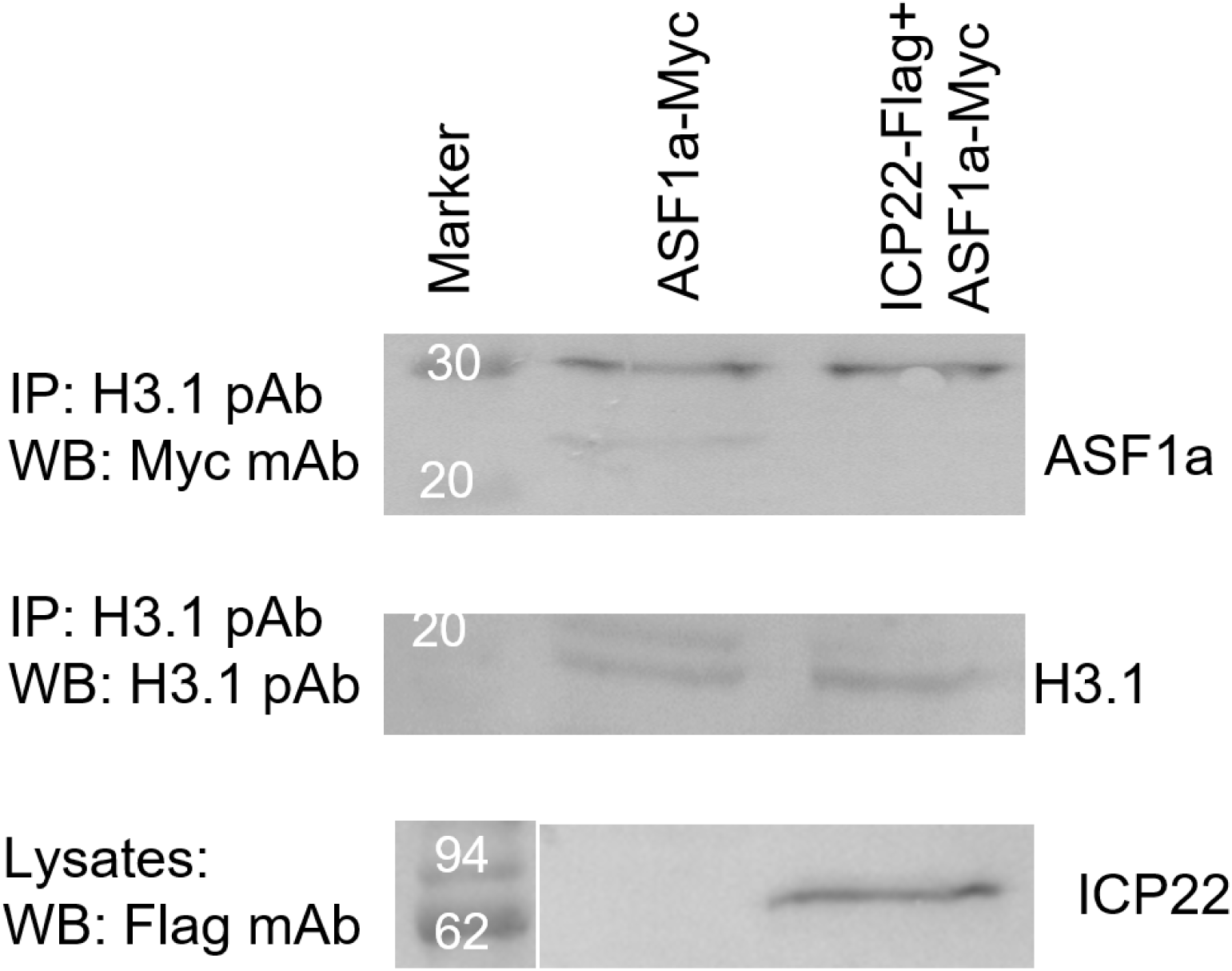
Ectopic expression of ICP22 abrogates the interaction between H3.1 and ASF1a. HEK293T cells were transfected with the pMyc-ASF1a plasmid or cotransfected with both the pMyc-ASF1a plasmid and the pcDNA-ICP22 plasmid. At 36 h after transfection, immunoprecipitation with a rabbit anti-H3.1 pAb and Western blot analysis were performed as described in Fig. 1A.

## DISCUSSION

ICP22 is a multifunctional HSV-1 regulatory protein that localizes predominantly to the nucleus of infected cells and has been linked to multiple transcription-associated processes during infection. Following the onset of DNA synthesis, ICP22 localizes to more diffuse transcription centers associated with L gene expression. Later in infection, ICP22 relocalizes together with the late UL3, UL4 and UL20.5 proteins to small dense nuclear bodies. In our previous study, the interaction relationships of the interaction complex among ICP22, UL3, UL4 and UL20.5 were comprehensively investigated [25]. Here, we identify ASF1 as an ICP22-associated host factor in cultured-cell systems and provide evidence that ICP22 associates with both ASF1a and ASF1b in transfected cells and HSV-1-infected cells.

ASF1 is an evolutionarily conserved histone H3/H4 chaperone involved in chromatin assembly, transcription-associated chromatin dynamics, and DNA replication-coupled histone management. ASF1, together with chromatin assembly factor (CAF1) and histones H3 and H4, assembles nucleosomes onto replicating DNA [31]. Additionally, ASF1 interacts with a wide range of chromatin-associated proteins, such as histone cell cycle regulation defective homolog A (HIRA) and TATA box binding protein-associated factor TAFII-250 [32], and with DNA damage checkpoint kinases, such as Tousled-like checkpoint kinase and Rad53 kinase [33]. Therefore, ASF1 has been implicated in various cellular functions, including DNA replication, gene transcription, and the cellular response to DNA damage, all of which involve orderly eviction and deposition of histones onto DNA [29]. Two mammalian isoforms, ASF1a and ASF1b, share a highly conserved central domain. The human ASF1a and ASF1b genes encode proteins of 204 amino acids (22.9 kDa) and 202 amino acids (22.4 kDa), respectively. They are highly conserved over much of their length but diverge in their amino and carboxyl termini. Both human ASF1a and human ASF1b bind the p60 subunit of CAF1 [34]. We found that HSV-1 ICP22 specifically interacts with both ASF1a and ASF1b in both transiently infected and HSV-1-infected cells.

Furthermore, amino acids 213 to 340 of ICP22 are responsible for its interaction with both ASF1a and ASF1b. Interestingly, amino acids 213 to 340 of ICP22 were among the conserved sequences in alphaherpesvirus. A previous study revealed that the VZV IE63 protein, the homolog of HSV-1 ICP22, also interacts with human ASF1 [20], suggesting that the conserved domains among alphaherpesvirus ICP22 homologous proteins are required for their interaction with ASF1. Additionally, the highly conserved aa 37 to 153 of ASF1a and ASF1b are required for their interactions with ICP22. The colocalization of ICP22 with ASF1a and ASF1b further demonstrated their interactions.

It has been previously reported that the cotransfection of ICP22 with a panel of HSV reporter genes results in the repression of reporter protein activity. Ectopic expression of ICP22 was demonstrated to repress the promoter activities of both ICP4-Luc and TK-Luc, which are HSV-1 promoters. However, cotransfection of ASF1a abolished the repression of ICP4 or TK promoter activity by ICP22, suggesting that ASF1 plays important roles in overcoming the general repression of transcription by ICP22. In addition, knockdown of both ASF1a and ASF1b significantly reduced HSV-1 replication, indicating that both ASF1a and ASF1b are required for effective HSV-1 replication. Interestingly, these results are consistent with a previous report that ASF1b is critical for effective HSV-1 viral DNA replication [35].

Histones are highly basic proteins that form complexes with DNA [36]. Ectopic expression of ICP22 was shown to abrogate the interaction between ASF1 and H3.1, suggesting that the interaction between ASF1 and H3.1 may play an important role in the regulatory functions of ICP22 in HSV-1 infection.

Several limitations of this study should be noted. First, the experiments were performed in cultured cell lines, and the relevance of these findings to primary epithelial cells, neurons, differentiated tissues, or senescent cell states remains to be determined. Second, although our interaction and reporter data support functional interplay between ICP22 and ASF1, additional mechanistic studies will be needed to define the causal sequence of events and to test their contribution in more physiologically relevant infection models.

In summary, we demonstrated that HSV-1 interacts with both ASF1a and ASF1b and identified regions of ASF1 and ICP22 that are important for their interaction. Additionally, both ASF1a and ASF1b are required for effective HSV-1 replication. Therefore, the ICP22–ASF1 interface may represent a candidate target for future studies aimed at disrupting chromatin-associated steps of HSV-1 infection.

## ACKNOWLEDGEMENT

We thank Drs. Peter D. Adams and Stephen Rice for their gifts, plasmids Myc-ASF1a and ASF1b; pcDNA-US1.5, pcDNA22-BA and pcDNA22-PS.

